# High throughput screening and identification of coagulopathic snake venom proteins and peptides using nanofractionation and proteomics approaches

**DOI:** 10.1101/780155

**Authors:** Julien Slagboom, Marija Mladić, Chunfang Xie, Freek Vonk, Govert W. Somsen, Nicholas R. Casewell, Jeroen Kool

## Abstract

Snakebite is a neglected tropical disease that results in a variety of systemic and local pathologies in envenomed victims and is responsible for around 138,000 deaths every year. Many snake venoms cause severe coagulopathy that makes victims vulnerable to suffering life-threating haemorrhage. The mechanisms of action of coagulopathic snake venom toxins are diverse and can result in both anticoagulant and procoagulant effects. However, because snake venoms consist of a mixture of numerous protein and peptide components, high throughput characterizations of specific target bioactives is challenging. In this study, we applied a combination of analytical and pharmacological methods to identify snake venom toxins from a wide diversity of snake species that perturb coagulation. To do so, we used a high-throughput screening approach consisting of a miniaturised plasma coagulation assay in combination with a venom nanofractionation approach. Twenty snake venoms were first separated using reversed-phase liquid chromatography, and a post-column split allowed a small fraction to be analyzed with mass spectrometry, while the larger fraction was collected and dispensed onto 384-well plates before direct analysis using a plasma coagulation assay. Our results demonstrate that many snake venoms simultaneously contain both procoagulant and anticoagulant bioactives that contribute to coagulopathy. In-depth identification analysis from seven medically-important venoms, via mass spectrometry and nanoLC-MS/MS, revealed that phospholipase A_2_ toxins are frequently identified in anticoagulant venom fractions, while serine protease and metalloproteinase toxins are often associated with procoagulant bioactivities. The nanofractionation and proteomics approach applied herein seems likely to be a valuable tool for the rational development of next-generation snakebite treatments by facilitating the rapid identification and fractionation of coagulopathic toxins, thereby enabling specific targeting of these toxins by new therapeutics such as monoclonal antibodies and small molecule inhibitors.

**Author summary:** Snakebite is a neglected tropical disease that results in more than 100,000 deaths every year. Haemotoxicity is one of the most common signs of systemic envenoming observed after snakebite, and many snake venoms cause severe impairment of the blood coagulation that makes victims vulnerable to suffering life-threating hemorrhage. In this study, we applied a combination of analytical and pharmacological methods to identify snake venom toxins from a wide diversity of snake species that interfere with blood coagulation. Twenty snake venoms were screened for their effects on the blood coagulation cascade and based on the initial results and the medical relevance of the species, seven venoms were selected for in-depth analysis of the responsible toxins using advanced identification techniques. Our findings reveal a number of anticoagulant toxins that have not yet been reported before as such. The methodology described herein not only enables the identification of both known and unknown toxins that cause impairment of the blood coagulation, but offers a throughput platform to effectively screen for inhibitory molecules relevant for the development of next generation snakebite treatments.

## 1. Introduction

Snakebite is a medically important neglected tropical disease, with up to 5.5 million people bitten annually (1). These bites result in as many as 1.8 million envenomings and 138,000 deaths each year, with three to five times that number of people said to suffer from long term morbidity (1–4). It is the rural poor agricultural workers (e.g. farmers, herdsmen, etc) of the tropics and sub-tropics who suffer the greatest burden of snakebite, with incidences and case fatality rates highest in south and south-east Asia and sub-Saharan Africa (1). In part this is due to socioeconomic reasons, as victims in these parts of the world often do not have rapid access to specialized medical care due to limited health and logistical infrastructure (5), which severely restricts access to snakebite therapy. The only specific therapy available for treating snake envenoming is antivenom. Antivenom comprises polyclonal antibodies, which are purified from the blood of horses or sheep immunised with small, sub-toxic, amounts of snake venom. A proportion of the resulting IgG antibodies are specific to the venom toxins used for immunisation, and thus rapidly neutralize their activity when these therapies are delivered intravenously to patients, particularly if treatment occurs soon after a bite. However, despite these products saving thousands of lives every year, they possess a number of technical limitations that ultimately restrict their clinical utility. Antivenoms exhibit limited paraspecific efficacy (with efficacy restricted mostly to the species used for immunisation), have poor dose efficacy (only 10-20% of antivenom antibodies are typically specific to the toxin immunogens) and exhibit high incidences of adverse effects (which can be as high as in 75% of cases) (6–8). Critically, these therapies are expensive, ranging from $50-350 per vial in Africa, for example (6). Often, treating a snakebite can require multiple vial (e.g. up to 10-20), which makes these therapies unaffordable to the majority of snakebite victims, and pushes those individuals further into poverty. The consequences are poor uptake of this product, weak demand, a lack of manufacturing incentive, and a cycle of undersupply of affordable efficacious antivenom to the regions where it is needed most (9). Consequently, the development of alternative low-cost, low dose, safe and paraspecifically efficacious antivenoms would be of great benefit to snakebite victims inhabiting impoverished regions of the world (10).

Snake venoms consist of a mixture of different protein and peptides that are used to kill, immobilise or incapacitate their prey (11). These ‘toxins’ vary at every taxonomic level and cause a range of different toxicities, including haemotoxic, neurotoxic and/or cytotoxic pathologies (12–14). The majority of snakebite deaths are thought to be caused by snakes whose venoms are predominately haemotoxic, usually resulting in haemorrhage and perturbations of the clotting cascade. Indeed, venom induced consumption coagulopathy (VICC) is said to be one of the most medically important pathologies caused by snakebite, as it leaves patients highly vulnerable to life-threatening haemorrhage (15). VICC is caused by the continual activation of the clotting pathway via procoagulant toxins present in snake venom, ultimately leading to a depletion of clotting factors (particularly fibrinogen) via their consumption, and incoagulable blood [4]. The majority of snake species that are known to cause VICC are vipers, however, certain Australian elapid snakes, and a small number of colubrid and natricine snakes from Africa and Asia have also been reported to possess procoagulant toxins capable of causing VICC in snakebite victims (7, 15).

Although there are a number of different toxins responsible for causing VICC, many are thought to be members of the snake venom serine protease (SVSPs) and snake venom metalloproteinase (SVMPs) toxin families (5). These toxins activate the blood clotting cascade via their activity on a variety of different clotting factors, of which several are found towards the end of the clotting cascade, including factor V, factor X and prothrombin (factor II). In the same venom, it is not uncommon for other toxins to directly target fibrinogen, resulting in either its degradation or the promotion of weak fibrin clots (7, 11, 15). In addition to procoagulant toxins, snake venoms have also been reported to contain proteins with anticoagulant properties, such as phospholipases A_2_s (PLA_2_s) and C-type lectin-like proteins (16–18). As mentioned earlier, venom variation is ubiquitous among snake species, and thus the different snake species that cause VICC are thought to have differing numbers of pro- and anti-coagulant toxins in their venom, which may target different components of the blood clotting cascade, even if the pathological outcome in a victim (e.g. VICC) is the same. This variation of venom constituents makes it particularly challenging to develop generic antivenom that could be used to treat snakebite across the world. Thus, identifying the key toxins from different medically-important snake species that are responsible for causing life-threatening coagulopathies would enable us to take an informed approach to generating new, targeted, therapies for snakebite.

Recently we developed a 384 well plate-reader based plasma coagulation assay (19), which gave us the opportunity to perform high-throughput profiling of haemotoxic snake venoms. In addition, we developed an ‘at-line’ nanofractionation setup for high-throughput venom screening of individual components in crude venoms towards selected bioactivities (20, 21). Briefly, crude snake venoms are first separated with liquid chromatography, followed by a post-column split, which gives the opportunity of mass spectrometry (MS) detection in parallel to high-resolution nanofractionation (i.e. 6s per well) onto 384-well plates for bioassaying. Subsequently, bioactivity chromatograms can be constructed by plotting the readout of each well against the time of the corresponding fraction, and when coupled to our coagulation assay, these chromatograms show peaks with positive (i.e. pro-coagulation) or negative (i.e. anti-coagulation) maxima for each bioactive compound (19). Because each bioactivity chromatogram has a corresponding MS chromatogram obtained in the parallel measurement, the accurate masses of the bioactive peaks can be determined when correlating the peaks from the bioassay chromatograms with the data obtained from MS. Thus, this approach enables us to separate snake venom into fractions, and identify the proteins present in those fractions that cause coagulopathic effects in the bioassay.

In this study, we selected venoms from 20 different snake species, covering a broad geographical distribution and different taxonomic families, known to interfere with the blood clotting cascade, and we screened them using the described nanofractionation approach to detect their pro- and anti-coagulant activities. Subsequently, we selected seven of these venoms for detailed characterization analysis whereby wells containing bioactive compounds were subjected to tryptic digestion for full protein identification via proteomics. Our methodological approach represents a rapid and straightforward method for identifying pro- and anti-coagulant toxins in snake venoms in a throughput manner. In line with the cumulative literature, our findings show that PLA_2_ toxins are frequently detected as anticoagulant bioactive toxins across the species tested, while serine proteases and metalloproteinases are common constituents of procoagulant venom fractions. Thus, this methodology facilitates the identification of both known and unknown coagulopathic toxins, and provides an amenable and targeted approach for testing the neutralisation of pathogenic snake venom toxins by next generation antivenoms.

## 2. Materials and Methods

### 2.1. Chemicals and stock solutions

The solvents and chemicals used in this research were of analytical grade. Acetonitrile (ACN) and formic acid (FA) were purchased from Biosolve (Valkenswaard, The Netherlands) and water was purified using a Milli-Q plus system (Millipore, Amsterdam, The Netherlands). Bovine plasma was obtained from Biowest (Amsterdam, The Netherlands). Iodoacetamide, β-mercaptoethanol, Argatroban, Ammonium bicarbonate and calcium chloride, were obtained from Sigma-Aldrich (Zwijndrecht, The Netherlands). Sequencing grade modified trypsin was purchased from Promega Benelux B.V. (Leiden, The Netherlands), and stored and handled according to the manufacturer’s instructions. The snake venom pools used in this study *Echis ocellatus* (Nigeria)*, Echis carinatus* (India)*, Echis carinatus* (UAE)*, Echis pyramidum leakeyi* (Kenya)*, Echis coloratus* (Egypt)*, Crotalus horridus* (USA)*, Macrovipera lebetina* (Uzbekistan)*, Daboia russelli* (Sri Lanka)*, Bothrops asper* (Costa Rica)*, Bothrops jararaca* (Brazil)*, Lachesis muta* (Brazil)*, Bothriechis schlegelii* (Costa Rica)*, Calloselasma rhodostoma* (captive bred, Thailand ancestry)*, Hypnale hypnale* (Sri Lanka)*, Trimeresurus albolabris* (Thailand)*, Trimeresurus stejnegeri* (Malaysia)*, Deinagkistrodon acutus* (China)*, Dispholidus typus* (South Africa)*, Rhabdophis subminiatus* (Hong Kong) and *Oxyuranus scutellatus* (Papua New Guinea). They were sourced in house from animals held in, or historical venoms stored in, the herpetarium at the Liverpool School of Tropical Medicine, UK. These species were selected based on previous reports of coagulopathic venom activity (7, 19, 22, 23). Venoms were in lyophilized form at 4°C, until reconstitution in water to make 5 mg/mL stock solutions, which were then aliquoted and stored at – 80°C until use.

### 2.2. Ethics statement

The maintenance and handling of venomous snakes was undertaken following approvals granted by the Animal Welfare and Ethical Review Board of the Liverpool School of Tropical Medicine and the UK Home Office (licence #X2OA6D134), and thus meet the national and legal ethical requirements set by the Government of the United Kingdom (under the Animal and scientific procedures act 1986) and EU directive 2010/63/EU.

### 2.3. Liquid chromatography, at-line nanofractionation and mass spectrometry

Liquid chromatography separation, parallel at-line nanofractionation and subsequent mass spectrometry analyses were performed in an automated fashion. A Shimadzu UPLC system (‘s Hertogenbosch, The Netherlands) was used for the LC separation. All the settings of the system were controlled with Shimadzu Lab Solutions software. Fifty microlitres of each sample was injected by a Shimadzu SIL-30AC autosampler, and the two Shimadzu LC-30AD pumps were set to a total flow rate of 500 µl/min. A 250×4.6 mm Waters Xbridge Peptide BEH300 C_18_ analytical column with a 3,5-μm particle size and a 300-Å pore size was used for separation of the venoms. The separations were performed at 30 °C in a Shimadzu CTD-30A column oven. Mobile phase A comprised of 98% H_2_O, 2% ACN and 0.1% FA, and mobile phase B comprised of 98% ACN, 2% H_2_O and 0.1% FA. A linear increase of mobile phase B from 0% to 50% in 20 min was followed by a linear increase from 50% to 90% B in 4 min and a 5 min isocratic separation at 90% B. The starting conditions (0% B) were reached linearly in 1 min and the column was then equilibrated for 10 min at 0% B. The column effluent was split post-column in a 1:9 volume ratio. The larger fraction was sent to a FractioMate^TM^ FRM100 nanofraction collector (SPARK-Holland & VU, Netherlands, Emmen & Amsterdam) or a modified Gilson 235P autosampler. The smaller fraction was sent to the Shimadzu SPD-M30A photodiode array detector followed by an Impact II QTOF mass spectrometer (Bruker Daltonics, Billerica, MA, USA). An electrospray ionization source (ESI) was equipped onto the mass spectrometer and operated in positive-ion mode. The ESI source parameters were: capillary voltage 4.5 kV, source temperature 180°C, nebulizer at 0.4 Bar and dry gas flow 4 L/min. MS spectra were recorded in the *m/z* 50–3000 range and 1 average spectrum was stored per s. Bruker Compass software was used for the instrument control and data analysis. LC fractions (1 per 6 s) were collected row by row in serpentine-like fashion on clear 384-well plates (Greiner Bio One, Alphen aan den Rijn, The Netherlands) using the in-house written software Ariadne or FractioMator when the Fractionmate was used, with a maximum of four plates in one sequence. Each chromatographic run was collected into 350 wells of one 384-well plate. After fractionation, the plates were evaporated overnight for approximately 16 h using a Christ Rotational Vacuum Concentrator RVC 2-33 CD plus (Salm en Kipp, Breukelen, The Netherlands). The plates were then stored at –20°C until further use, i.e., either bioassaying or tryptic digestion.

### 2.4. Plasma coagulation assay

The coagulation cascade interfering properties of snake venoms were analysed using a previously described plasma coagulation assay (19). Briefly, 20 µL of 20 mM calcium chloride solution (room temperature) was dispensed onto a 384-well plate containing LC fractionated snake venom using a Thermo Scientific multidrop 384 reagent dispenser. Next, 20 µL of bovine plasma (room temperature) was dispensed into each well. Finally, the plate was placed into the plate reader and the absorbance was measured kinetically at 595 nM for 80 cycles (± 90 min) at 25°C.

The resulting data was normalized and analysed in three different ways to differentiate between anticoagulant, fast procoagulant and slow procoagulant detected bioactive components First, the last measurement cycle was plotted as a single point measurement to display the anticoagulant compounds. Second, the average slope of cycles 1-5 were plotted to show the fast procoagulant components. Last, the average slope of cycles 1-15 were plotted to show the remainder of the procoagulant components (“slow procoagulants”). The rationale for plotting the procoagulant activity in two different ways is that the fast procoagulant compounds clot the plasma in the wells in a distinct manner, resulting in a rapid rise in absorbance followed by a plateau, and where this plateau typically occurs at a lower overall absorbance than that observed with the “slow procoagulant” components. Thus, wells containing the slower procoagulant components reach an absorbance that surpasses those of the wells containing the fast procoagulants, but over a prolonged period of time.

### 2.5. Correlating coagulation data with mass spectrometry data

The elution times and peak widths at half maximum of the peaks observed in the bioassay chromatograms were determined. For the corresponding time interval, the average mass spectrum was extracted from the parallel recorded MS chromatogram (total-ion current (TIC)). For all m/z’s with a significant signal in this mass spectrum, extracted-ion currents (XICs) were plotted. Based on matching peak shape and retention time, m/z’s were assigned to the appropriate bioactive peaks observed in the bioassay chromatogram based. The charge states of the particular one deconvolutes spectra were assigned by the software based on the observed isotope distribution. Finally, deconvolution of the full mass spectra recorded for the assigned peaks provided the accurate monoisotopic masses.

The difference in flow rate of the flow ratio directed to the bioassay and the flow ratio directed to the mass spectrometer after the post-column split caused a constant time delay between the nanofractionated plates and the obtained MS data. To determine the duration of this delay, the thrombin inhibitor argatroban was nanofractionated, resulting in a negative peak in the plasma coagulation assay, which could be correlated to its corresponding known extracted ion current (shown in SI Fig 1). The difference in elution time between the two peaks is the delay between the MS and the nanofractionation collector and was found to be 78 sec. Using the determined delay, the *m/z*-values measured in the MS could be correlated with the bioactive peaks observed in the bioassay chromatograms.

### 2.6. Identification of coagulopathic toxins via tryptic digestion

After for a specific sample the wells containing bioactive compounds were pinpointed, the same sample was fractionated again, and the content of the active wells was subjected to tryptic digestion in order to identify coagulopathic toxins using a proteomics approach. For that, the plate was first freeze-dried, and then 40 µL of water was added to each selected well, reconstituting the contents for 30 min at room temperature. Next, 10 µL from each rwell was transferred to an Eppendorf tube containing 15 µL of digestion buffer (25 mM ammonium bicarbonate, pH 8.2). Subsequently, 1.5 µl of reducing agent (0.5% β-mercaptoethanol) was added to each tube followed by a 10-min incubation at 95 °C. Then, the samples were cooled to room temperature and centrifuged at 150 RCF for 10 s in a Himac CT 15RE centrifuge. Next, 3 µL of alkylating agent (55 mM iodoacetamide) was added and the mixture was incubated in the dark at room temperature for 30 min. Subsequently, 3 µL of 0.1 μg/μL trypsin was added followed by incubation at 37 °C for 3 h, after which an additional 3 µL of trypsin solution was added to the tube and the samples were incubated overnight. Next, 1 μL of 5% FA was added to quench the digestion, followed by brief centrifugation to remove any particulate matter. Finally, the samples were centrifuged for 10 s at 150 RCF and the supernatants were transferred to autosampler vials with glass inserts and analyzed using nanoLC-MS/MS.

NanoLC separation of the tryptic digests were performed using an UltiMate 3000 RSLCnano system (Thermo Fisher Scientific, Ermelo, The Netherlands). The autosampler was run in full-loop injection mode. The autosampler was set to a 1 µL injection volume and after injection the samples were separated on an analytical capillary column (150 mm x 75 µm) packed in-house with Aqua C_18_ particles (3 µm particle size, 200 Å pores; Phenomenex, Utrecht, The Netherlands). The mobile phase was comprised of eluent A (98% water, 2% ACN, 0.1% FA) and eluent B (98% ACN, 2% water, 0.1% FA). The applied gradient was: 2 min isocratic separation at 5% B, linear increase to 80% B in 15 min, 3 min isocratic separation at 80% B, down to 5% B in 0.5 min and equilibration for 9 min. The column was kept at 30 °C in the column oven. Absorbance detection was performed at 254 nm followed by mass detection using a Bruker MaxisII qTOF mass spectrometer (Bruker, Bremen, Germany) equipped with a nanoESI source operating in positive-ion mode. The ESI source parameters were: source temperature, 200 °C; capillary voltage, 4.5 kV; gas flow, 10 L/min. Spectral data were stored at a rate of 1 average spectrum/s in the range of 50 to 3000 m/z. MS/MS spectra were obtained using collision induced dissociation (CID) in data-dependent mode using 35-eV collision energy. Bruker Compass software was used for instrument control and data analysis.

Using the data obtained for the tryptic digests of the samples, protein identification was carried out using MASCOT (Matrix Science, London, United Kingdom) searches against Swiss-Prot, NCBInr and species-specific databases. The latter were generated from previously published transcriptomic data for those species that were available (24–27) and were used for the identification of venom toxins for which activity was observed. The following search parameters were used: ESI-QUAD-TOF as the instrument type, semiTrypsin as the digestion enzyme allowing for one missed cleavage, carbamidomethyl on cysteine as a fixed modification, amidation (protein C-terminus) and oxidation on methionine as variable modifications, ± 0.05 Da fragment mass tolerance and ± 0.2 Da peptide mass tolerance. To compare the deconvoluted monoisotopic masses with masses found by Mascot we used Uniprot to obtain information about potential PTMs and the presence or absence of pro and signal peptides. Uniprot contains transcriptomic data which usually does not represent the native form of the toxins since it does not take possible PTMs and the absence of pro or signal peptides into account. Therefore toxins were drawn in Chemdraw including the appropriate PTMs and excluding any pro or signal peptides in order to obtain the exact mass to facilitate correlations between XICs and the MS data. The Uniprot database generated a high number of Mascot identities from which a selection was made based on protein score (> 50), sequence coverage (> 20%) and known activity.

### 2.7. Statistical analysis

Comparisons of the molecular weights of the various procoagulant and anticoagulant venom toxins detected from the seven venoms were performed using the R package Artool (28). The data comprised of the molecular weights of toxin identities found through tryptic digestion and Mascot database searching described above. The full data set of toxin molecular weights can be found in Table 2. As the compiled data was found to be non-Gaussian in distribution, a non-parametric factorial analysis was performed, which was undertaken using the function art() in Artool (28).

## 3. Results & Discussion

### 3.1. Coagulopathic activity profiling of snake venoms

The objective of this study was to characterise the coagulopathic activity of various snake venoms and identify their coagulopathic toxins. Our approach consisted of an initial screening strategy, whereby the bioactivity of 20 snake venoms sourced from different taxonomic families and geographic origins were elucidated, followed by in depth proteomic characterization of the active coagulopathic compounds for seven of those venoms. The initial HT screening approach utilized high-resolution LC fractionation of venom (50 µL injections of 5 mg/mL solutions) into a 384-well plate coupled to a plasma coagulation assay. This bioassay measures differences in clotting velocity between control wells which produce a normal clotting profile (e.g. plasma in the presence of calcium chloride), and wells containing bioactive components (i.e. procoagulants or anticoagulants). Plotting the different velocities against the time of the fraction results in positive peaks for procoagulant compounds (where clotting is more rapidly stimulated, resulting in enhanced clotting speed) and negative peaks for anticoagulant compounds (where clotting is reduced or inhibited, resulting in prolonged clotting times or prevention of clotting).

Based on the initial screening, the pro- and anti-coagulant activities of the 20 snake venoms were compared semi-quantitatively based on the observed activity (Table 1). For a detailed view of the resulting assay data for each of these venoms we direct the reader to SI Fig 2. Our findings demonstrate a variable pattern of coagulopathic activity among those species at the concentration tested. The majority of the venoms tested demonstrated procoagulant activities, which was anticipated given prior findings (7), although the extent of this activity varied extensively, with *Bothrops asper* providing the most potent effect. Surprisingly, neither *Oxyuranus scutellatus* or *Rhabdophis subminiatus* venoms exhibited evidence of procoagulation under the assay conditions used here, despite prior reports of this venom activity (7, 22). Similarly, we observed extensive variation in anticoagulant venom effects, with three of the 20 species showing maximal effects (Table 1), while we did not detect discernible anticoagulant venom activity for nine of the species. Notably, *B. asper* also exhibited the most potent anticoagulant effects detected (alongside *Daboia russelii* and *Oxyuranus scutellatus*), demonstrating that this venom is highly coagulopathic, and acts via both procoagulant and anticoagulant activities.

**Table 1.** Semi-quantitative determination of procoagulant activity of venom of 20 snake species. The extent of pro- and anti-coagulant activity is graded from strong (+++) to weak (+), where 0 represents no activity detected.

| Species | Pro | Anti | Species | Pro | Anti |
| --- | --- | --- | --- | --- | --- |
| *Echis ocellatus* | + | 0 | *Bothriechis schlegelii* | ++ | ++ |
| *Echis carinatus* | ++ | ++ | *Calloselasma rhodostoma* | ++ | 0 |
| *Echis carinatus* | + | 0 | *Hypnale hypnale* | ++ | 0 |
| *Echis pyramidum leakeyi* | + | ++ | *Trimeresurus albolabris* | + | + |
| *Echis coloratus* | + | ++ | *Trimeresurus stejnegeri* | + | ++ |
| *Crotalus horridus* | + | 0 | *Deinagkistrodon acutus* | + | 0 |
| *Macrovipera lebetina* | + | ++ | *Dispholidus typus* | + | 0 |
| *Daboia russelii* | ++ | +++ | *Rhabdophis subminiatus* | 0 | 0 |
| *Bothrops asper* | +++ | +++ | *Oxyuranus scutellatus* | 0 | +++ |
| *Bothrops jararaca* | ++ | 0 | *Lachesis muta* | ++ | ++ |

Seven venoms (underlined in Table 1) were selected for in depth characterisation. These venoms were selected because they represent snakes of greatest medical importance in their distinct geographic locales (e.g. the viperid species *Echis ocellatus*, *Daboia russelii*, *Bothrops asper*, *B. jararaca*, *Calloselasma rhodostoma*) or because they represent different snake families (e.g. the elapid species *Oxyuranus scutellatus* and the colubrid species *Dispholidus typus*). The results of these optimized experiments demonstrated that all seven of the snake venoms exhibited substantial procoagulant venom activity (Figure 1), broadly consistent with our initial venom screen (Table 1), and recent analyses undertaken using crude venom (7, 22, 29–35). The notable exception to this was that of *O. scutellatus*, which did not exhibit procoagulant activity in our initial screen (Table 1), but did so in these optimized experiments (Figure 1). These latter findings are more consistent with the literature than our earlier experiment (22), and likely reflect the differences in venom concentration (5mg/mL instead of 1mg/mL) and CaCl_2_ (20 mM instead of 100 mM) used for the subsequent optimized experiments. The decrease of CaCl_2_ resulted in an assay readout with significantly less baseline noise and therefore increased sensitivity, which combined with increased venom concentration, resulted in the detection of procoagulant venom activity *for O. scutellatus*.

**Fig 1.**
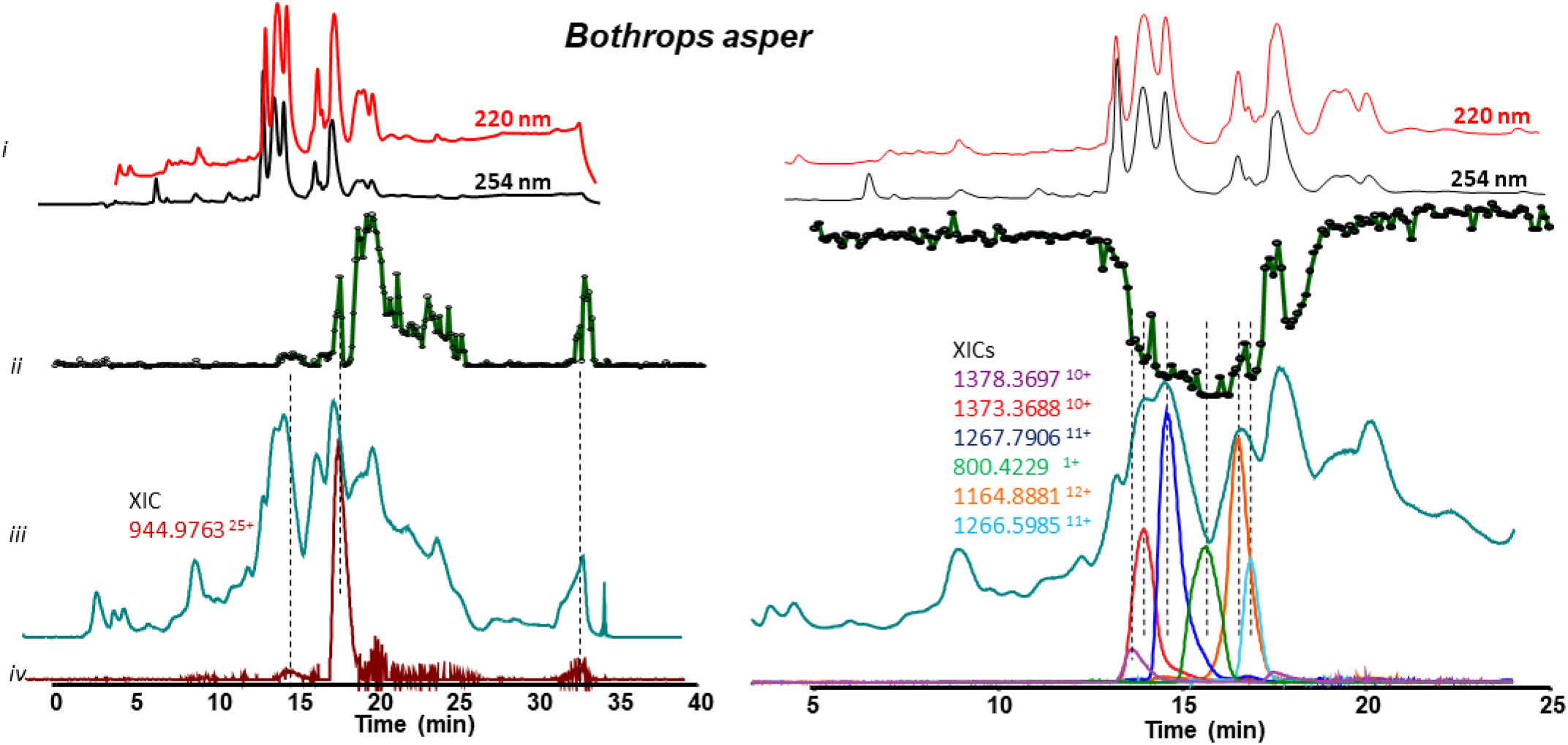
Detection of coagulation interfering compounds of *Bothrops asper* by correlating MS and bioassay data obtained upon LC analysis of crude snake venom (5 mg/mL). *i*: UV chromatograms detected at 220 and 254 nm. *ii*: Bioactivity chromatograms obtained after fractionation by plotting the results of the plasma coagulation assay against time; the positive and negative peaks indicate the presence of procoagulant and anticoagulant bioactive compounds respectively. *iii*: MS chromatograms (TIC), *iv*: Extracted-ion chromatograms (XICs) of *m/z*-values corresponding to bioactive peaks. Further experimental conditions, see Materials and Methods section.

### 3.2. Proteomic identification of coagulopathic toxins

#### 3.2.1. Correlation of bioactivity peaks with mass spectrometry data

After coagulopathic profiling of the venoms, the first step in identifying the active toxins was to assign accurate masses of individual venom components exhibiting activity in the plasma coagulation assay. This was done by correlating bioactivity chromatograms to the corresponding MS chromatograms. This correlation was based on elution time and peak shape of the bioactive peaks. The data interpretation of the analysis of the seven snake venoms assayed here are shown in figures 1 and 2, and the *m/z*-values and their corresponding masses for all bioactives can be found in table 2. This method proved to be successful for masses up to 15 kDa, however larger toxins showed to be much more difficult to correlate. This was most likely caused by a lack of sensitivity of the LC-MS caused by poor ionization or insufficient amounts of the relevant proteins.

**Fig 2.**
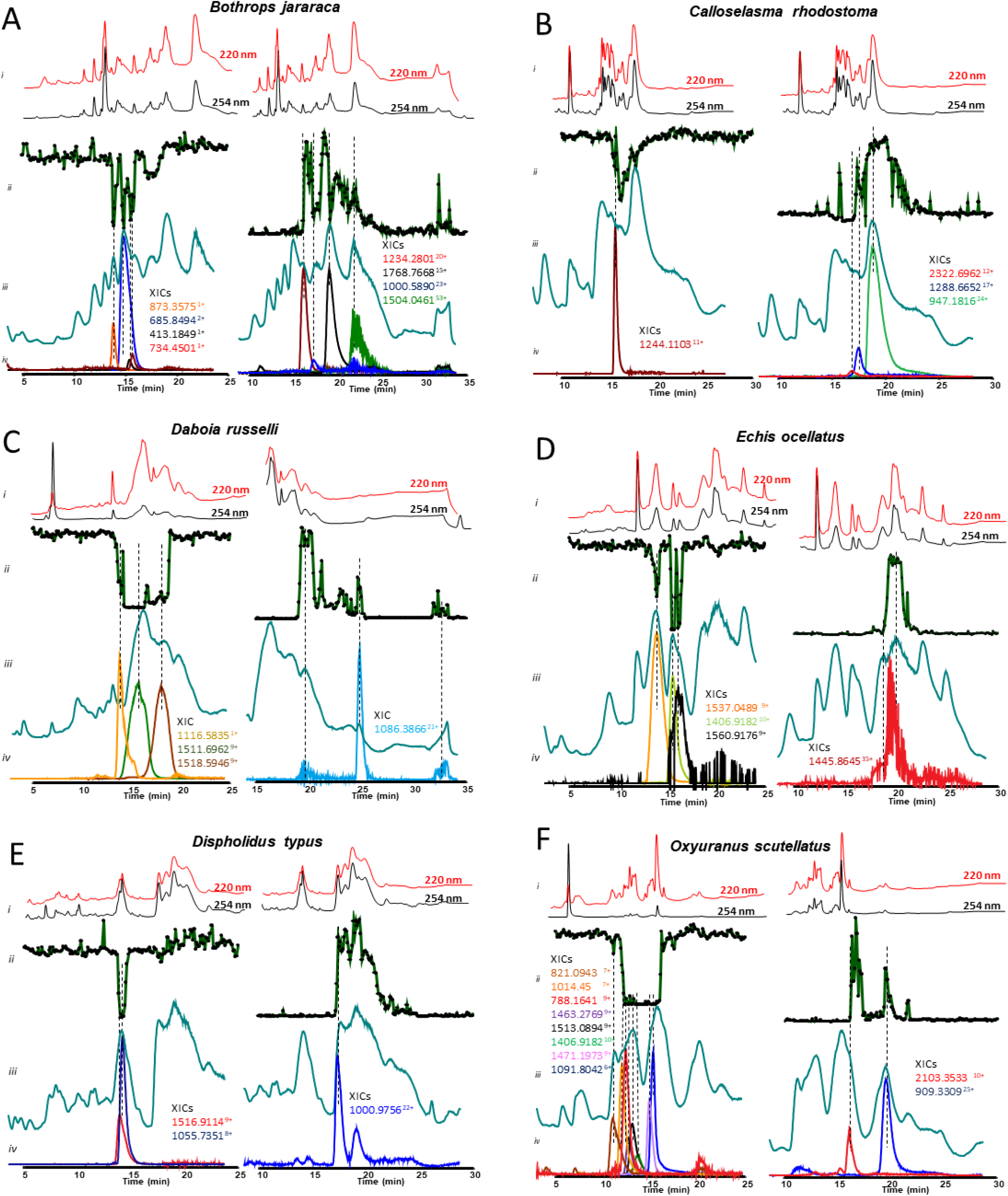
Identification of coagulopathic compounds from the remaining six venoms by correlating MS data with bioassay data. *i*: UV trace of the snake venoms at 220 and 254 nm obtained by LC-MS. *ii*: bioactivity chromatograms obtained by plotting the results of the plasma coagulation assay in Prism software. The peaks with positive (left) and negative (right) maxima indicate the presence of procoagulant and anticoagulant bioactive compounds respectively. 6-s resolution fractions were collected onto 384 well plates by the nanofractionator after 50-µL injection of crude snake venom at a concentration of 5 mg/mL. *iii*: LC-MS chromatograms displaying the total ion current (TIC), *iv*: the extracted ion currents (XICs) of the *m/z*-values corresponding to bioactive peaks found in the MS spectra.

**Table 2.**
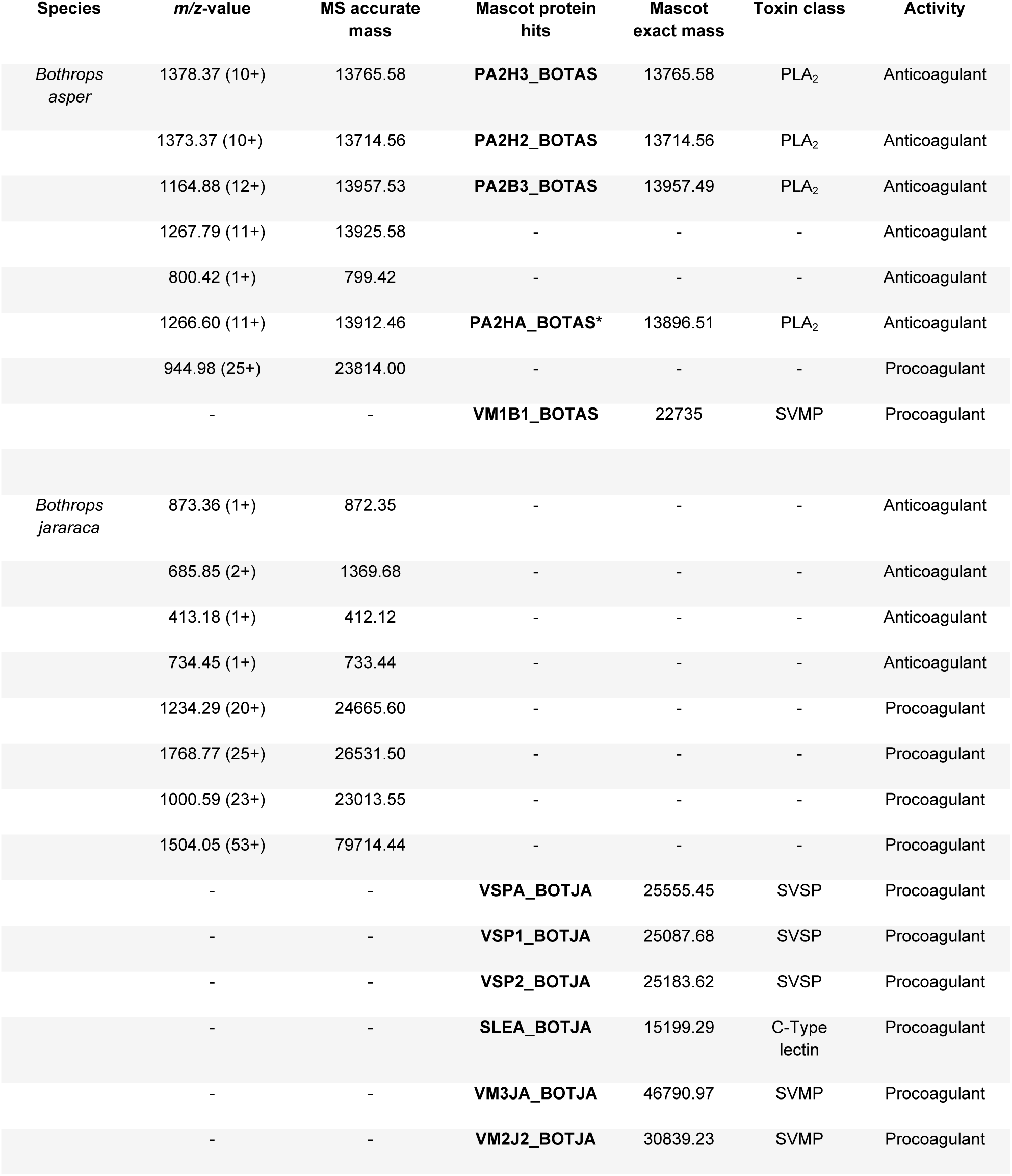
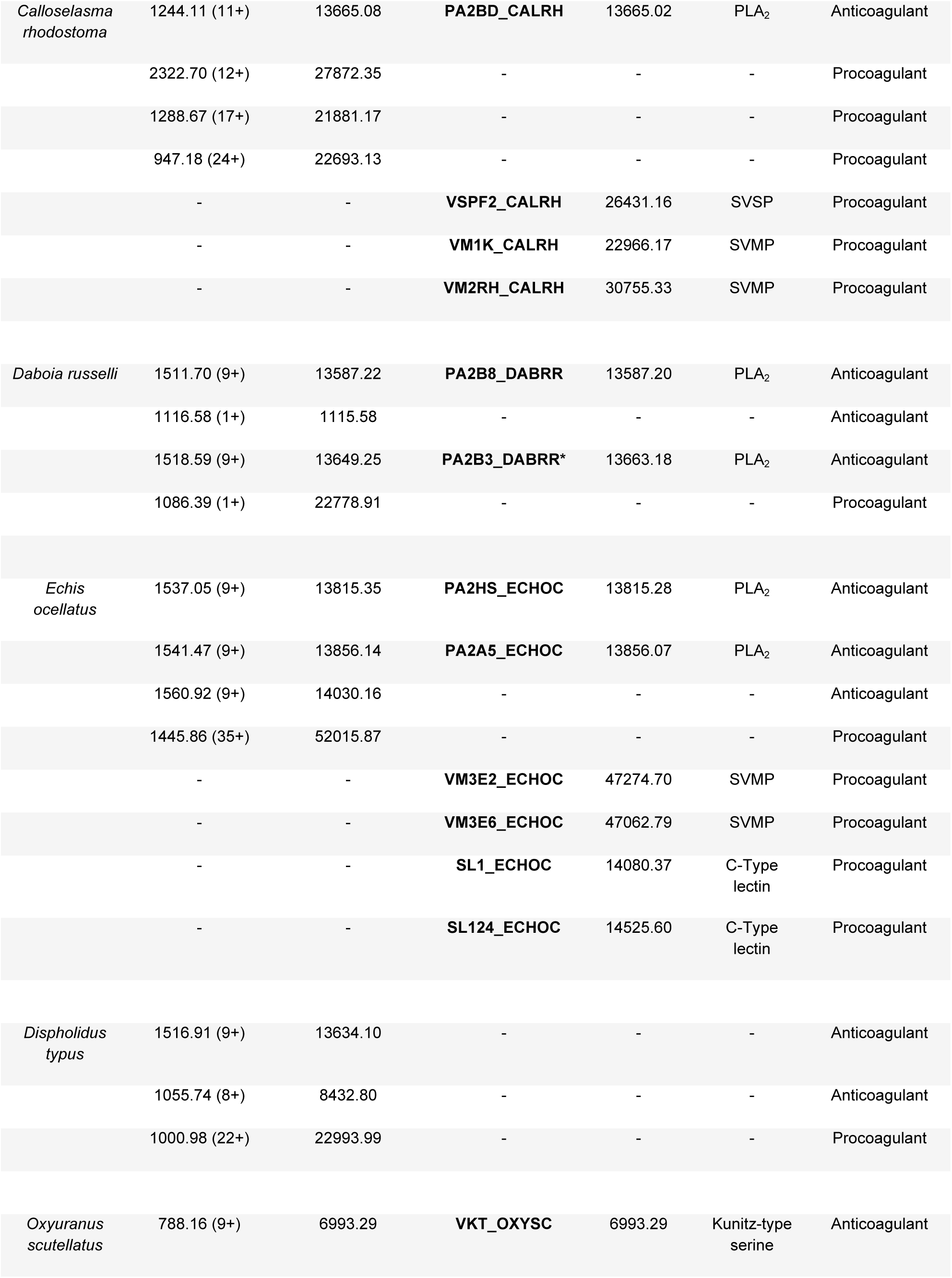
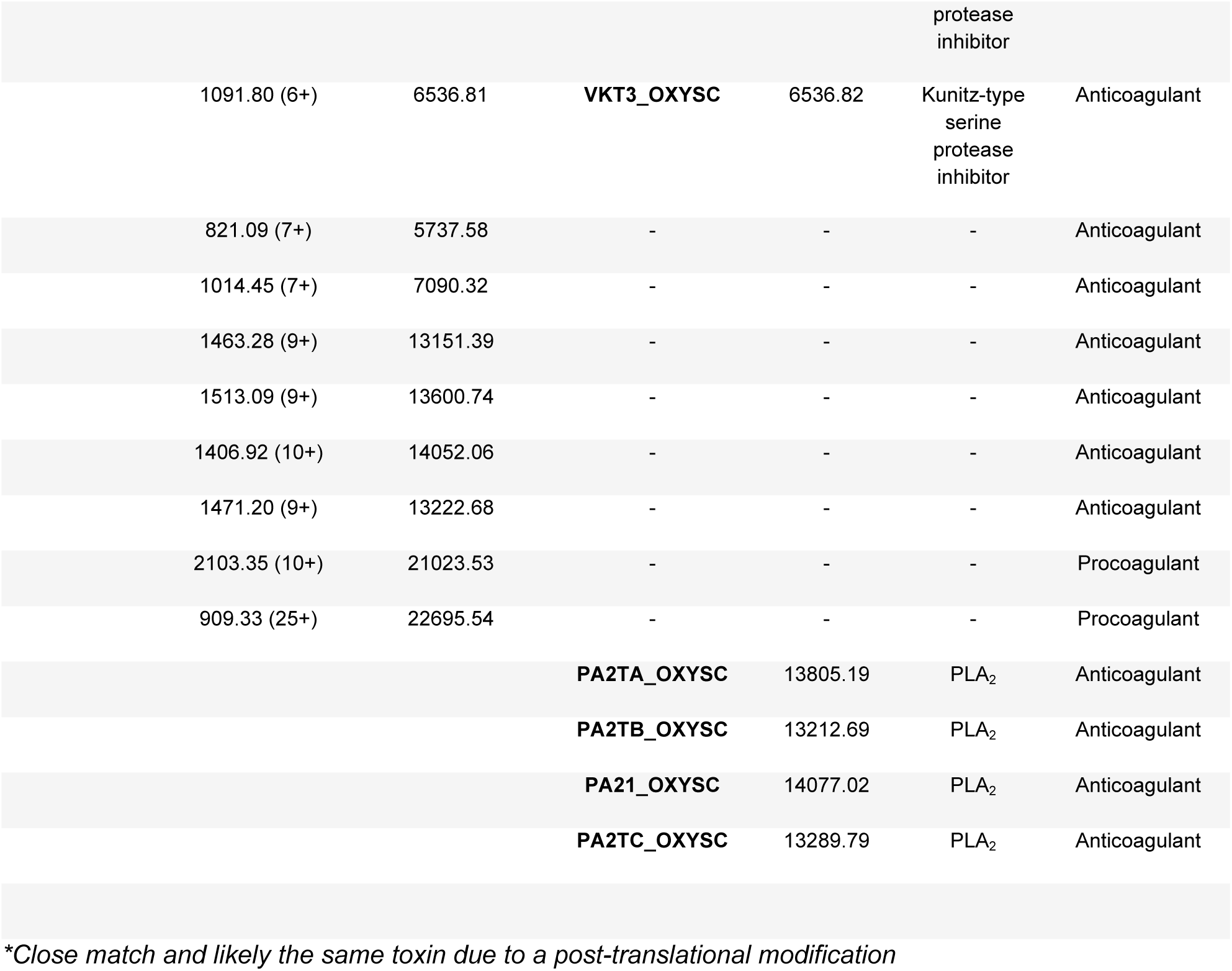
Assignment of coagulopathic venom toxins based on masses detected for intact bioactive components and Mascot hits after tryptic digestion.

#### 3.2.2. NanoLC-MS/MS analysis of tryptic digests of venom fractions showing bioactivity

In the second step to identify the coagulopathic venom toxins observed in the plasma coagulation assay after fractionation, venoms were re-fractionated and the content of selected wells containing bioactive compounds was subjected to tryptic digestion (see SI Fig 3 for details of well selection for each venom). The digested content of each well was analysed by nanoLC-MS/MS and the data was subjected to a Mascot database search using the Uniprot database and species-specific databases compiled from transcriptomic data. We noted that the search results provided by the Uniprot database provided a large number of Mascot identities at the timeframes of the pro- and anti-coagulant activities. This is likely the result of the bioactive compound not having an exact match in the database due to low toxin representation, but having high sequence similarity to a number of venom toxins found in related snake species. The toxin identities mentioned in the text below were selected based on the sequence coverage, protein score and known functional activity, and the full results are detailed in the SI Uniprot database table in the Supplementary Information. Thereafter, we compared the deconvoluted monoisotopic masses of the XICs correlated with the bioactive peaks (described above) with the masses of the compounds identified via Mascot to assign protein identities found on Uniprot to coagulopathic venom toxins (Table 2). This proved to be again successful for smaller toxin components (<15 kDa), but for larger toxins (e.g. >20kDa) none of the XICs could be correlated to Mascot identities (Table 2).

Despite these limitations, summarizing the toxin identities from the Mascot hits and XICs of the various anti- and pro-coagulant venom fractions revealed noticeable differences in the masses of the toxins associated with these two bioactivities (Fig 3. Panel C). In total we detected 59 toxins in the fractions where anticoagulant activities were observed for the seven venoms. The masses ranged from 412 Da to 30 kDa in size, with the majority falling between 1 and 15 kDa. For procoagulant activity we found 38 toxins, which exhibited a much broader mass range, from 11 kDa to 79 kDa. However, the majority of these were larger than 15 kDa in size. Although there are clear inter-specific distributions in the molecular weights of the various coagulopathic toxins identified, statistical analyses reveal that procoagulant venom toxins exhibit significantly higher molecular weights than their anticoagulant counterparts (non-parametric factorial analysis; *P* = 2.22e-16, F=163.52). In the next sections we detail the toxin identities of the various coagulopathic toxins found in each of the seven venoms.

**Fig 3.**
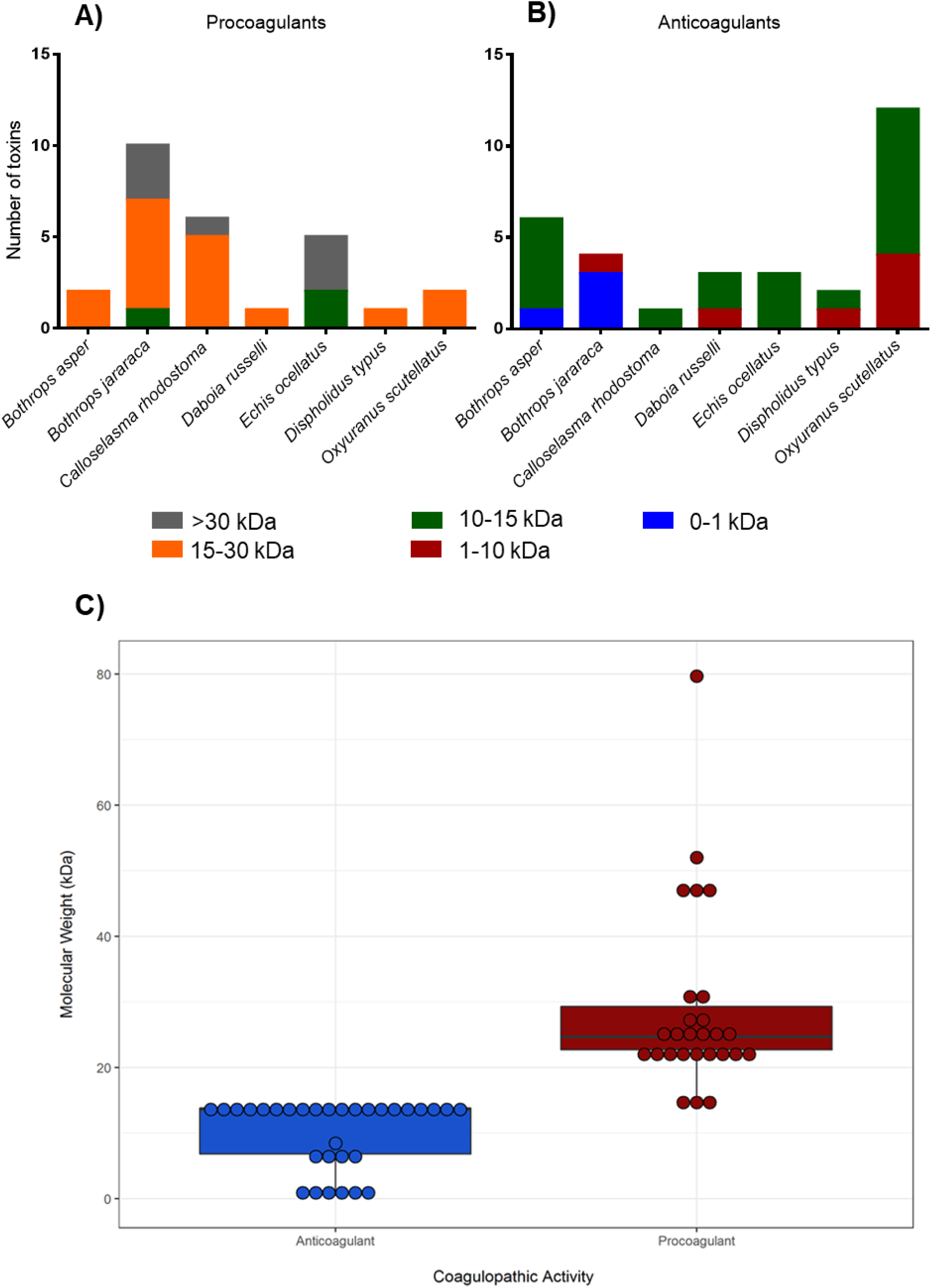
The molecular weight distribution of the various anti- and pro-coagulant toxins identified in the venom of each of the seven snake species. The number and molecular weight range of toxins identified from each snake species are summarized for procoagulant (**A**) and anticoagulant (**B**) bioactives. Toxin identifications are derived from those described in Table 2 and their respective molecular weights were calculated by drawing structures in chemdraw and adding appropriate PTMs. The full data can be found in detail in the SI Uniprot database table. **C**) Molecular weight comparisons of the anticoagulant and procoagulant toxins detected from all seven snake venoms. Boxes show the interquartile range of each dataset, with the boldened horizontal line representing the median value, and each data point is illustrated by a dot. Statistical comparisons, via non-parametric factorial analysis, reveals that procoagulant venom toxins exhibit significantly higher molecular weights than anticoagulants (*P* = 2.22e-16, F=163.52).

### 3.3. Mascot results summary for all seven species

#### 3.3.1 Bothrops asper

On the basis of the *m/z*-values detected for the anticoagulant-active peaks for *Bothrops asper*, six molecular masses were assigned (Figure 1; Table 1). Four of these were found to be proteins of the PLA_2_ toxin family. A number of snake venom PLA_2_s have previously been described to exert anticoagulant effects via binding to factor Xa and inhibiting prothrombinase activity (36). One of the four PLA_2_s detected in our study has previously been described to be anticoagulant, namely “Basic phospholipase A2 homolog 2” (*m/z*-value of 1373.3688^10+^, which corresponds to an accurate mass of 13714.5639 Da), which has been described to inhibit prothrombinase activity (36). The remaining three PLA_2_s have *m/z*-values of 1378.3697^10+^, 1266.5985^+11^ and 1164.8811^12+^, which correspond to accurate masses 13765.5821 Da, 13912.4649 Da and 13957.5232 Da, respectively. The first of these identities corresponds to the “PLA_2_ Basic phospholipase A2 homolog M1-3-3”, which is predicted to exert myotoxic activity based on its similarity to PLA_2_s from other snake species (37). The second corresponds to “Basic phospholipase A2 homolog 4a” and exhibits a mass ∼16 Da higher than the PLA_2_ toxin, and this slight difference may be caused by an oxidation of methionine. This toxin has only previously been described as myotoxic (38). The third PLA_2_ corresponds to “Basic phospholipase A2 myotoxin III” which is also only known to exhibit myotoxic activity (38–43). The remaining masses detected to potentially exert anticoagulant activity had an XICs of 800.4229^1+^ and 1267.79^11+^ and accurate masses of 799.4155 and 13925.58 Da, which both remain unidentified. Despite *B. asper* venom exhibiting potent procoagulant activity (e.g. Table 1, Figure 2) only one *m/z*-value could be assigned to components exerting this effect, 944.976325^+25^, and to which no corresponding match could be found in the Mascot database after proteomics analysis. Nonetheless, given that the mass of this venom toxin was 23.8 kDa, it seem likely that it belongs to the SVMP or SVSP toxin family. Analysis of Mascot identities found at the time frame of the eluted procoagulant activities, detected relatively few toxins, but of those “Snake venom metalloproteinase BaP1” is a known procoagulant SVMP, and thus may be responsible for this observed bioactivity (41).

#### 3.3.2 Bothrops jararaca

Despite the association of eight *m/*z-values with coagulopathic venom activity, we were unable to confidently assign any of these venom component masses to Mascot hits. Four of these components corresponded to anticoagulant bioactive peaks in the plasma coagulation assay (Figure 2B), and exhibited masses of 400-1400 Da. These findings contrast starkly with the majority of the anticoagulants identified in the venom of the congeneric species *B. asper* (five anticoagulants 12-14 kDa in size), with two unidentified components from that species (799 and 13925.58 Da) with 799 Da falling within a similar mass range to the anticoagulants identified in *B. jararaca* venom. While some of these low molecular weight components may simply be co-eluting with anticoagulant toxins, it is also possible that the active toxins are indeed co-eluting but are not detected by either MS or Mascot. However, further elucidation of these compounds could be of particular value if they do exhibit anticoagulant activity since the small size of these molecules makes them amenable from a drug discovery perspective. Bradykinin-potentiating peptides match the detected molecular masses and, have previously been characterized for this venom (44, 45). As for *B. asper*, none of the m/z-values associated with procoagulant activity could be matched to Mascot hits. Three of these components exhibited masses in the 23-27 kDa range, whereas the fourth had a mass of about 80 kDa (Table 2). The Mascot hits found at the timeframe where the procoagulant activity was observed suggest that some of these proteins may be SVSPs that are known to exhibit procoagulant (“Thrombin-like enzyme bothrombin”; sequence coverage (SC) of 22%), platelet aggregating (“Platelet-aggregating proteinase PA-BJ”; SC, 49%) or fibrinogenolytic (“Thrombin-like enzyme KN-BJ 2”; SC, 40%) activities. In addition, we detected matches to C-type lectins with known platelet aggregating activities (“Snaclec botrocetin subunit alpha”; SC, 20%) (46), and SVMPs previously described to promote haemorrhage and haemagglutination (“Zinc metalloproteinase-disintegrin-like jararhagin” and “Zinc metalloproteinase/disintegrin”; SC, 46% and 28%, respectively) (40, 46–49).

#### 3.3.3 Calloselasma rhodostoma

In total, four bioactive coagulopathic toxins were detected in the venom of *C. rhodostoma*, one that corresponded to an anticoagulant bioactivity peak, and three to procoagulant peaks (Figure 2C). The anticoagulant correlated to the XIC 1244.1103^11+^ and had a resulting accurate mass of 13665.0848 Da. When comparing this mass with the Mascot search results, this venom toxin was matched to “Inactive basic phospholipase A2 W6D49”, a PLA_2_ toxin known to cause local oedema but has yet to be described as having anticoagulant activity (50). Similar to those venoms previously described, the XICs could not be correlated to procoagulant bioactivity peaks or Mascot identities. However, the masses of these procoagulant venom toxins were 21-23 kDa and ∼28 kDa in size, which suggests that they likely belong to the SVMP or SVSP toxin families. Indeed, the Mascot hits found where procoagulant activity was observed revealed the presence of both these toxin types, along with C-type lectins and an L-amino acid oxidase (Table 2). The SVSP identified (“Thrombin-like enzyme ancrod-2”; SC, 53%) has previously been shown to be fibrinogenolytic (51), while the literature relating to the SVMPs detected (“Snake venom metalloproteinase kistomin” and “Zinc metalloproteinase/disintegrin” SC, 33% and 27%, respectively) suggests they predominately exert anticoagulant activities, including interfering with platelet aggregation and promoting haemorrhage (52, 53). It is therefore possible that these toxins are co-eluting with procoagulant components, although given that many SVMPs are multi-functional proteins that can exert procoagulant bioactivities (54), they may be contributing to the functional effect observed here despite these activities not previously being reported.

#### 3.3.4 Daboia russelii

The venom of *D. russelii* exhibited both potent anticoagulant and procoagulant bioactivities (Figure 2D). We identified three XICs that correlated to anticoagulant bioactivity peaks, one of which exhibited a *m/z*-value (1511.6962^9+^) and could be directly matched to a Mascot identity. This protein had an average mass of 13587.2248 Da and corresponded with “Basic phospholipase A2 VRV-PL-VIIIa” (sequence coverage of 82%) - a PLA_2_ toxin that has been previously described to be anticoagulant (36). Of the remaining anticoagulant XICs, one exhibited a mass of 13649.2513 Da (*m/z*-value 1518.5946^9+^) and could be correlated to “Basic phospholipase A2 3” (55) exhibiting a mass ∼14 Da higher than the PLA2 toxin, and this slight difference may be caused by a methylation., whereas the other protein was much smaller in size (*m/z*-value 1116.5835^1+^; average mass 1115.5759 Da), and remains unidentified. We identified one XIC (*m/z*-value 1086.3866^21+^) that correlated with a bioactive procoagulant peak, but were also unable to match this via Mascot database searches. The molecular mass of this venom toxin was 22778.9073 Da, which raises the possibly of it being an SVMP or SVSP toxin.

#### 3.3.5 Echis ocellatus

We identified three anticoagulant bioactivity peaks in the venom of *Echis ocellatus* that could be linked to XICs (Figure 2E), and two of these were matched to Mascot hits. Both of these XICs (*m/*z-values of 1537.0489^9+^ and 1541.4718^9+^) resulted in similar average mass values (13815.3523 Da and 13856.1382) and matched the PLA_2_ toxins “Phospholipase A2 homolog” and “Acidic phospholipase A2 5”. Despite these identifications, and prior reports of PLA_2_s generally exerting anticoagulant activities (16), neither of these *E. ocellatus* PLA_2_s have previously been described as anticoagulants (33, 56, 57). Despite the venom from these species being potently procoagulant (7, 58), we only identified one XIC that could be correlated to a procoagulant bioactivity peak. This protein had a *m/z*-value of 1445.8645^35+^ and a corresponding mass of 52015.873 Da, but could not be matched to a Mascot hit. Nonetheless, the mass recovered suggests that this could be a SVMP, particularly since this venom is known to be dominated by this toxin class (33), and has previously been demonstrated to contain P-III SVMPs of ∼52 kDa in size (13), and potent procoagaulant SVMPs, such as the ∼55 kDa prothrombin activator ecarin (59). Supporting this tentative toxin assignment, Mascot hits at the timeframe of the procoagulant activity identified known procoagulant SVMP toxins (“Zinc metalloproteinase-disintegrin-like EoVMP2” and “Zinc metalloproteinase-disintegrin-like EoMP06” SC, 48% and 23% sequence coverage, respectively) (60, 61), along with two, much small molecular weight, C-type lectins (“Snaclec 1” and “Snaclec CTL-Eoc124” SC, 53% and 31%). “Zinc metalloproteinase-disintegrin-like EoVMP2” is a known prothrombin activator and “Zinc metalloproteinase-disintegrin-like EoMP06” is suspected to activate prothrombin based on its sequence similarity with a prothrombin-activating SVMP of *E. pyramidum leakyi* (Ecarin, A55796, 91% similarity) (60, 61). While it is known that some SVMPs form complexes with CTLs, including the *Echis* prothrombin activator carinactivase (62), it is unlikely that these complexes would remain intact after nanofractionation.

#### 3.3.6 Dispholidus typus

The venom of *D. typus* yielded three potential coagulopathic venom toxins, two of which were associated with anticoagulant activity, and the third with procoagulanct effects. Two m/z-values (1516.9114^9+^ and 1055.7351^8+^; masses, 13634.0965 and 8432.8015 Da) were correlated with the anticoagulants. However, these two venom toxins are closely eluting (Figure 2F) and thus it is possible that only one of these components is responsible for the anticoagulant activity. Based on their molecular weight it is tempting to speculate that this activity likely comes from the ∼13 kDa compound, particularly since this mass correlates well with a number of anticoagulants detected from the other venoms under study here (e.g. PLA_2_ toxins, 13-14 kDa in size). However, given that *D. typus* is the only colubrid venom investigated in this study, and that colubrid PLA_2_s are distinct from those of vipers (24), this hypothesis requires testing in future work. Nonetheless, these findings are interesting because there are no prior reports of *D. typus* venom exhibiting anticoagulant activity, likely because this activity is masked by the net procoagulant activity of this venom (7). The procoagulant peak could be correlated to a mass of ∼23 kDa (Table 2). This toxin most likely belongs to the SVMPs family, which are the most dominant toxin type in *D. typus* venom (24), and have previously been described to be responsible for the potent procoagulant activity of this species (63).

#### 3.3.7 Oxyuranus scutellatus

*Oxyuranus scutellatus* is the sole representative of the elapid snake family under study here. As venom toxin composition diverted following the split of vipers from all other caenophidian snakes around 50 million years ago (64), we anticipated identifying a number of toxins types not found in viper venoms. The bioactivity chromatogram revealed a large elution region of anticoagulants, with no distinct peaks (Figure 2F). The m/z-values of eight compounds could be correlated to this anticoagulant region. Two of these (788.1641^9+^ and 1091.8042^6+^; masses, 6993.2858 and 6536.8065 Da) could be matched via Mascot and were identified as Kunitz-type serine protease inhibitors (Kunitz-type serine protease inhibitor taicotoxin and Kunitz-type serine protease inhibitor scutellin-3) both of which are known to exhibit anticoagulant activities and are abundant components in *O. scutellatus* venom (22, 27, 35). The remaining m/z-values had no associated Mascot match. Two of these unidentified components had molecular masses of 5.5-7.1 kDa and thus may also be anticoagulant Kunitz toxins. However, the remaining four anticoagulant XICs corresponded to molecular masses in the range of ∼13-14 kDa, and therefore seem likely to be PLA_2_s, particularly since the venom of *O. scutellatus* is rich in this toxin class (22). This assertion is also supported by four known PLA_2_s being detected in the timeframe of the observed anticoagulant activity, namely: Basic phospholipase A2 taipoxin alpha chain, Neutral phospholipase A2 homolog taipoxin beta chain 1, Phospholipase A2 OS1 and Neutral phospholipase A2 homolog taipoxin beta chain 2. However, none of these PLA_2_s have previously been described to be anticoagulant (65, 66). Despite this, and bearing in mind that elapid PLA_2_s are distinct from those of vipers (67), a number of elapid PLA_2_s have been demonstrated to exert anticoagulant effects (68, 69). Nonetheless, improved separation of these venom toxins, followed by isolation and retesting, is required to confirm these tentative identifications. Two m/z-values were correlated to procoagulants with molecular masses of 21-23 kDa, but neither could be matched to Mascot results, and thus remain unidentified from this study.

## 4. Study limitations

Summarising across the species studied here, we found that many of the anticoagulants detected are likely PLA_2_ toxins, as indicated by matching masses for some, and the molecular weights determined by MS for others (i.e. ∼13-14 kDa). Contrastingly, only a few of the procoagulant activity peaks observed in the plasma coagulation assay could be assigned to detected masses, and none were identified via Mascot database searches. Most probably, these components are of relatively large molecular mass and are poorly ionized, and transferred under the applied ESI and MS conditions (70). Using more dedicated conditions for intact protein analysis would likely overcome this limitation (71, 72). Masses of compounds that could be correlated to procoagulants ranged from 20 to 80 kDa, with the majority between 21 and 28 kDa. Compounds in this mass range most likely are SVMPs and SVSPs, which contain toxins that have previously been described to exhibit procoagulant activities in many haemotoxic snake venoms (11, 18, 73).

Although venom toxin enzymes are known to be relatively stable under typical RPLC separation conditions (i.e. exposure to organic solvent), on-column denaturation can occur, particularly in the case of later eluting venom toxins. Moreover, in RPLC, proteins typically yield rather broad peaks (in contrast to peptides) and representatives of multiple toxin families might elute within the same retention time frames due to poor resolution. Examples observed from this study include PLA_2_s coeluting with C-type lectins, and SVMPs with SVSPs. Thus, for some of the potential coagulopathic toxins observed, the assignments described earlier are necessarily tentative, as assignment to specific proteins is hindered by co-elution.

The nanoLC-MS analysis of the tryptic digests of bioactive fractions followed by Mascot database searches did, however, provide a large number of hits, indicating that venom toxins are present but were not previously detected as intact proteins directly after separation by MS. More details regarding the Mascot SwissProt database search is found in the SI. This includes Mascot finding toxins in the seven analysed snakes from different species, which is explained by the fact that there are high sequence similarities for many toxins among different snake species. From all the Mascot data obtained two extensive tables were comprised, one generated from the Uniprot database and the other from the species-specific databases (see SI Species specific databases table and SI Uniprot databases table). There was a significant difference in recovering components above 20 kDa between the LC-MS-bioassay correlation approach and the nanoLC-MS approach. This was most likely caused due to the lack of sensitivity of the LC-MS caused by poor ionization or insufficient amounts of the proteins, as mentioned before. The nanoLC-MS-Mascot approach was far more sensitive due to the much better ionization of peptides compared to intact proteins. Isolation and further elucidation of the toxins correlated to the bioactivity peaks is necessary for the confirmation of their activity. Finally, it is possible that there could be co-eluting bioactive components that were not detected by MS.

It has to be mentioned that for most of the seven snakes profiled many more Mascot identities were found at the timeframes of the pro- and anticoagulant activities but that mostly only the ones for which a high sequence coverage and/or an accurate mass was established, were mentioned in the text. The venom toxin identities reported are in some cases from different species than analyzed, which means that the possible bioactive compound is not an exact match, but has high sequence similarity to a venom toxin from another snake species of which the transcriptomic data was available in the Mascot database. The sequence similarities between species are causing a high number of search hits in Mascot and this is the reason why not all findings are mentioned in the text but presented in the SI Uniprot databases table.

## 5. Summary of toxin identifications

Despite these limitations, the at-line nanofractionation approach in combination with the plasma coagulation assay proved to be a fast and powerful approach for the identification of a number of venom toxins that affect the coagulation cascade. Fig 4., provides a summary of the toxins that were detected as being procoagulant and anticoagulant from each of the seven snake venoms studied. For the procoagulant activities, SVMPs, SVSPs and C-type lectins were found in five out of the seven species analysed in depth, while LAAOs were found in three of the seven (Fig 4). These identifications form the basis for future confirmatory studies characterizing the procoagulant activities of the toxins identified here, and for defining those venom components that to not perturb coagulation and were detected here as the result of co-elution. For the anticoagulant activities it became evident that PLA_2_s play an important role for all species included in this study, as at least one PLA_2_ was detected in each of the seven venoms. In six of the seven venoms analysed we also detected C-type lectins in bioactive regions, although only one of these was associated with anticoagulant activity (in *Oxyuranus scutellatus*). This is perhaps surprising given that C-type lectins are predominately associated with anticoagulant activities (17, 74–78), however, these toxins may perhaps work in a synergistic manner with other toxins, or may not have been associated with anticoagulant activity here, due to many eluting in the procoagulant timeframe of the bioassay, resulting in anticoagulant activities being masked by procoagulant toxins. While the reverse may also happen (e.g. weakly procoagulant toxins being masked via co-eluting in the anticoagulant areas), it is worth noting that one of the strengths of our approach is the deconvolution of coagulopathic toxin activities from the complex venom mixture, and specifically the identification of anticoagulant venom activity from these ‘net’ procoagulant venoms. Prior studies using crude venom have demonstrated that the majority of the venoms tested here are procoagulant and potently clot plasma (7, 29, 31, 32, 54, 79–82). Despite these observations, our nanofractionation approach clearly reveals that many of these venoms also contain anticoagulant bioactives (e.g. 10 of 20 initial venoms, Table 1; 7 of 7 venoms analysed in depth; Figures 1 and 2), which are completely masked when using crude venom in such screening assays.

**Fig 4.**
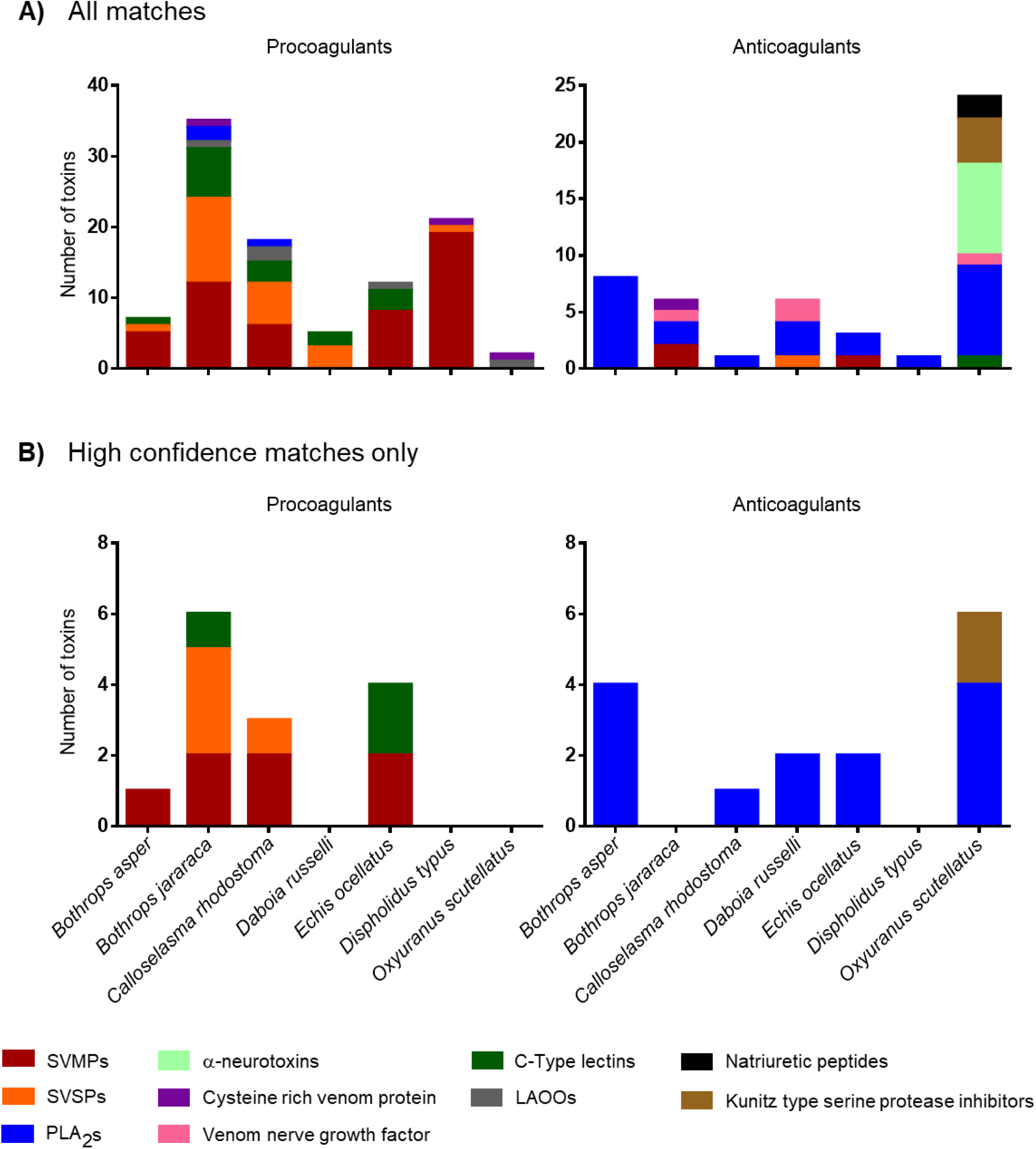
Identification of the various anti- and pro-coagulant toxins associated with the areas in the bioactivity profiles where pro- and anti-coagulant activities were observed. Panel A shows all the toxins found where pro and anticoagulant is observed. This panel is constructed with the SI species specific databases table found in the Supplementary information. Panel B shows toxins that could be matched to observed bioactivity with high confidence due to correlation with MS data or abundance of the toxin and its known activity. This panel is constructed with the data found in Table 2.

## 6. Conclusion

The results of this study describe the validity of using a medium throughput approach for the identification of venom toxins associated with specific pathogenically-relevant bioactivities. In this case, we applied the screening approach to the topic of coagulopathy, as this is well known to be one of the most frequent, serious, pathologies observed following snakebite. The mechanisms of action of coagulopathic snake venom toxins are diverse and can result in both anticoagulant and procoagulant effects. Characterisation of specific target bioactives is challenging though, as venoms are complex mixtures of toxins, therefore methods to deconvolute them are required. The methodology applied here facilitated the rapid identification and fractionation of coagulopathic toxins, which resulted in the identification of anticoagulant venom activities previously being masked by net procoagulant effects of the crude venom, and also detected a number of anticoagulant PLA_2_s not previously known to exhibit anticoagulant activities. Success here, alongside the recent development of other small-scale medium-throughput bioassays focused on characterizing other relevant toxin activities (20, 83–86) means that such assays could be used interchangeably for understanding venom function and pathology – ultimately providing a ‘bioassay toolkit’ for deconvoluting venom mixtures. These approaches will undoubtedly facilitate new research focused on improving snakebite therapy. Current conventional antivenoms are hampered by their low specificity, as they consist of polyclonal antibodies generated via the immunization of animals with snake venom. This process results in only 10-20% of antibodies being specific to venom toxins, with the remainder directed against environmental antigens, and of those specific antibodies only a proportion will actually be specific to the key pathogenic toxins found in any particular venom (87). Thus, much research effort is now focusing upon increasing the specificity of therapies to specific, pathogenically-important, venom toxins, including the novel application of small molecule inhibitors and monoclonal antibodies (88–93). For both these approaches, screening and identification platforms are important for the selection of molecules exhibiting desirable *in vitro* neutralizing profiles from large drug or antibodies libraries. Thus, ultimately, we foresee the application of this methodological platform to greatly enable the future development of new snakebite therapies, by providing a rational screening process that will enable targeted testing of venom toxin neutralization by new inhibitory molecules.

## 7. Acknowledgements

The authors thank Taline Kazandjian for assistance with statistical analyses, and Paul Rowley for the extraction of snake venoms. This study was supported by: (i) a Sir Henry Dale Fellowship to N.R.C. (200517/Z/16/Z) jointly funded by the Wellcome Trust and Royal Society, and (iii) a UK Medical Research Council funded Research Grant (MR/S00016X/1) to N.R.C. and J.K.

## 9. Supporting Information Legends

**SI.** SI Fig 1. Correlation of mass spectrometry data with assay data using argatroban, SI Fig 2. Initial screening results of all 20 species included in the study, SI Fig 3. Toxin IDs from the specific and non-specific databases from nanofractionated toxins present in wells and subjected to proteomics.

**SI_Species specific databases table.** All Mascot hits found with the species specific databases in the area where pro- and anticoagulant activity was observed. Table includes information on masses, retention times, well numbers of the nanofractionated toxins, sequence coverage, protein score, toxin class and coagulation activity.

**SI_Uniprot database table.** All Mascot hits found with the Uniprot database in the area where pro- and anticoagulant activity was observed. Table includes information on masses, retention times, well numbers of the nanofractionated toxins, sequence coverage, protein score, toxin class and coagulation activity.

